# Antibody-conjugating nanogel (Conjugel) with two immune checkpoint inhibitors for enhanced cancer immunotherapy

**DOI:** 10.1101/2023.10.19.563185

**Authors:** Yun Jin Chae, Kang-Gon Lee, Doogie Oh, Su-Kyoung Lee, Yongdoo Park, Jongseong Kim

## Abstract

Cancer immunotherapy by immune checkpoint inhibitors (ICI) acts on antitumor responses by stimulating the immune system to attack cancer cells. However, this powerful therapy is hampered by its high treatment cost and limited efficacy. Here, we show the development of an antibody-conjugating system (Conjugel) that potentiates the efficacy of bispecific immunotherapy that simultaneously targets CTLA-4 and PD-L1. The Conjugel, consisting of highly deformable nanogels and antibody-binding protein, was loaded with two ICI monoclonal antibodies (mAb). Compared with mAb treatment alone, treatment with a bispecific Conjugel loaded with the both ICIs significantly decreased both the survival of MCF-7 and MDA-MB-231 breast cancer cells *in vitro* and the size of 4T1-Luc2-derived orthotopic syngeneic tumors *in vivo*. Furthermore, the ICI-loaded Conjugel was less toxic *in vivo* than the combination treatment delivered as a bolus. Our findings have important implications for Conjugel-based immunotherapy, developing the safer and higher efficacy of ICIs to treat breast cancers.

## Introduction

Immunotherapy using immune checkpoint inhibitors (ICIs) is a powerful strategy in the treatment of multiple cancer types^1–6^. To date, two types of ICIs, characterized by their ability to block the immunomodulatory receptor cytotoxic T lymphocyte-associated antigen 4 (CTLA-4) and the immunoinhibitory receptor programmed death 1 (PD-1) and its ligand, programmed death ligand 1 (PD-L1), have been approved by the FDA and clinically administered for the treatment of solid tumors, including breast cancers^7^. Although the use of immunotherapy in clinical oncology has shown remarkable success in cancer treatment over the past decade, the limitations of cancer immunotherapy, such as its high cost and low patient response rate, have also become more pronounced, which hints at un-met needs for maximizing its therapeutic potential ^1, 7–11^.

One attempt to address these restrictions involves the use of combination therapy using two kinds of ICIs or an ICI with another cancer drug^5, 12–17^. Such combination therapy has been shown to be more efficacious than monotherapy in a number of clinical studies^6, 18, 19^, but associated adverse effects lead to discontinuation of this therapy^1, 15, 20, 21^. Colitis and myocarditis are the most common causes of deaths in patients treated by ICI-related combination therapies^21, 22^. Furthermore, another method to address the limitations of ICIs involves the development of bispecific or multispecific antibodies (Abs) that can simultaneously target two or more antigens, respectively, through genetic engineering and chemical conjugation^23–25^. In cancer immunotherapy, such antibodies are designed to both activate T lymphocytes and target cancer cells to provide greater therapeutic effects, similar to combination therapy^26–28^. However, the clinical use of these antibodies is hampered by not only the time-consuming and high-cost nature of their production but also the hyperimmunogenicity of artificial antibodies.

As another potential method to improve ICI efficacy, the use of drug delivery systems (DDSs) in cancer immunotherapy is a novel approach to not only extend the application of ICIs to patients who have not benefited from this powerful therapy but also improve efficacy for better clinical outcomes^28, 29^. This is because advanced DDSs allow the delivery of cancer drugs to a targeted region over a specific time frame, improving therapeutic potential and decreasing adverse effects^30^. In particular, among DDSs, nanoparticles offer several advantages; for instance, they protect therapeutic agents from physiological inactivation and enable their accumulation in the targeted tissue^30–34^. Cell-based DDS technologies, such as chimeric antigen receptor (CAR) T-cell therapy, have also been developed to recruit therapeutic molecules to living cells and aim to reprogram native immune cells to attack cancer cells^35^. Despite the revolutionary success of CAR-T cell therapies, their high cost and complex and time-consuming production restrain their extensive use in cancer treatment^10^. Thus, advancing cell-like platforms for use in DDSs to overcome the aforementioned hurdles is desirable.

Here, we describe a novel linking and delivery platform for cancer immunotherapy, an Antibody-Antibody Conjugel (AAC). The advantage of this platform over current engineered bispecific antibodies and DDS modalities arises from their adaptability to link multiple antibodies quickly and easily, enabling AAC system to be a novel platform for the rapid development of enhanced therapeutic formulation in cancer treatment. This also help prevent those antibodies from randomly contacting and killing noncancerous cells, where subcutaneously injected into tumor region. We tested those features of AAC in two types of breast cancer cells (MCF-7 and MDA-MB-231) *in vitro* and orthotopic syngeneic model (4T1-Luc2) *in vivo*.

## Results

### Viability assays of breast cancer cells

It has been reported that blocking CTLA-4 promotes the activation of cytotoxic T cells via CD28 pathways, while inhibition of PD-L1 eliminates the ability of cancer cells to evade the immune system^36,37^. To implement those therapeutic effects in our system, we devised an Antibody-Antibody Conjugel (AAC) that consists of highly deformable hydrogel nanoparticles loaded with one or two ICIs – anti-CTLA-4 and anti-PD-L1 (Fig. 1a). Poly(*N*-isopropylacrylamide-*co*-acrylic acid) (pNIPAm-AAc) nanogels synthesized by the precipitation polymerization method were embedded by protein A through a carbodiimide coupling method via EDC/NHS ^38^. The Fc regions of the ICIs (monoclonal antibodies (mAbs) were specifically attached to the protein A-modified nanogels, allowing exposure of their Fab regions and, in turn, binding to the corresponding target molecules, i.e. CTLA-4 and PD-L1 (Fig. 1b). After binding, the ICIs remain attached to the protein A modified nanogels, i.e. Conjugel. Brightfield and fluorescence microscopy showed not only that the ICI-conjugated AAC were successfully formed (Fig. 1c), but also that binding of the anti-PD-L1-conjugated AACs to the MCF-7 cancer cells was significantly greater than that of the isotype control. (Fig. 1d, e).

**Figure 1.**
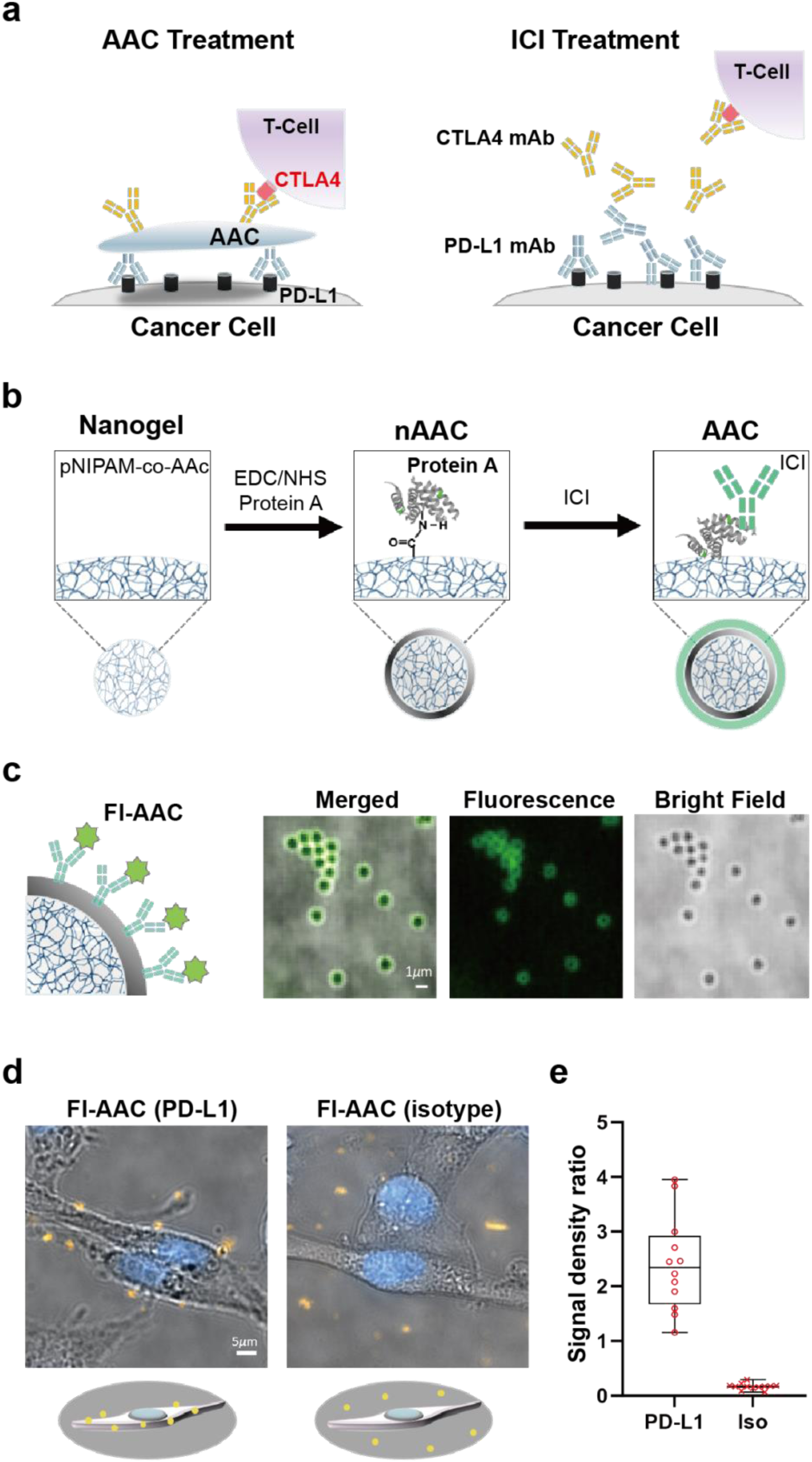
Antibody-antibody Conjugel (AAC)-based cancer immunotherapy and AAC preparation for cancer cell targeting. **a**, Schematic diagrams of the bispecific AAC equipped with CTLA-4 mAb and PD-L1 mAb and its interaction with cancer cells in the presence of peripheral blood mononuclear cells (PBMCs). **b,** The poly(*N*-isopropylacrylamide-*co*-acrylic acid) (pNIPAm-AAc nanogels (left) is conjugated with protein A (nAAC) via EDC/NHS coupling (center), forming AACs complexed with the Fc regions of the ICIs (right). **c,** Fluorescence and bright field microscopy images representing the formation of the PD-L1 mAb (ICI)-conjugated AAC. AAC was incubated with fluorescently labeled PD-L1 mAb (Fl-AAC), which was then attached to an amine-functionalized cover glass for imaging with a 100X oil immersion objective lens. The scale bar represents 1 μm. **d-e,** Targeting of the PD-L1 mAb-conjugated AAC to cancer cells. Fluorescence and bright field microscopy images showing the localization of PD-L1 or the isotype mAb-conjugated Fl-AAC in MCF-7 breast cancer cells, respectively (**d**), and analysis of the fluorescence signal density ratio between the cell surface and the bottom of the cover glass (**e**). The scale bar represents 5 μm. Significance between PD-L1:Fl-AAC and isotype:Fl-AAC was assessed using the Mann‒Whitney U test. Note that 11 spot areas of PD-L1:Fl-AAC and 22 spot areas of isotype:Fl-AAC were assessed for analysis.

To evaluate the efficacy and toxicity of the AAC to cancer cells and normal cells, we designed an *in vitro* assay in which both kinds of cells were cocultured with peripheral blood mononuclear cells (PBMCs) and treated with each AAC or ICI alone. More specifically, two types of breast cancer cells (cell lines MCF-7 and MDA-MB-231) and two types of normal cells (cell lines MCF-10 and AC16) were each treated with three types of AAC containing one or both of the ICIs examined in the study (anti-PD-L1:AAC, anti-CTLA-4:AAC, and anti-PD-L1:anti-CTLA-4:AAC) and three ICI treatment regimens (single anti-PD-L1 treatment, single anti-CTLA-4 treatment, and combination anti-PD-L1:anti-CTLA-4 treatment) (Fig. 2a). We first confirmed that the PBMCs and the AAC without an ICI (nAAC) had a nonspecific effect on the viability of the two types of breast cancer cells (Fig. S1). Monospecific AACs with either anti-PD-L1 or anti-CTLA-4 (mono-AACs) were selected to target cancer cells or immune cells, respectively, and a bispecific AAC with anti-PD-L1 and anti-CTLA-4 (bi-AAC) was used to simultaneously modulate both types of cells.

**Figure 2.**
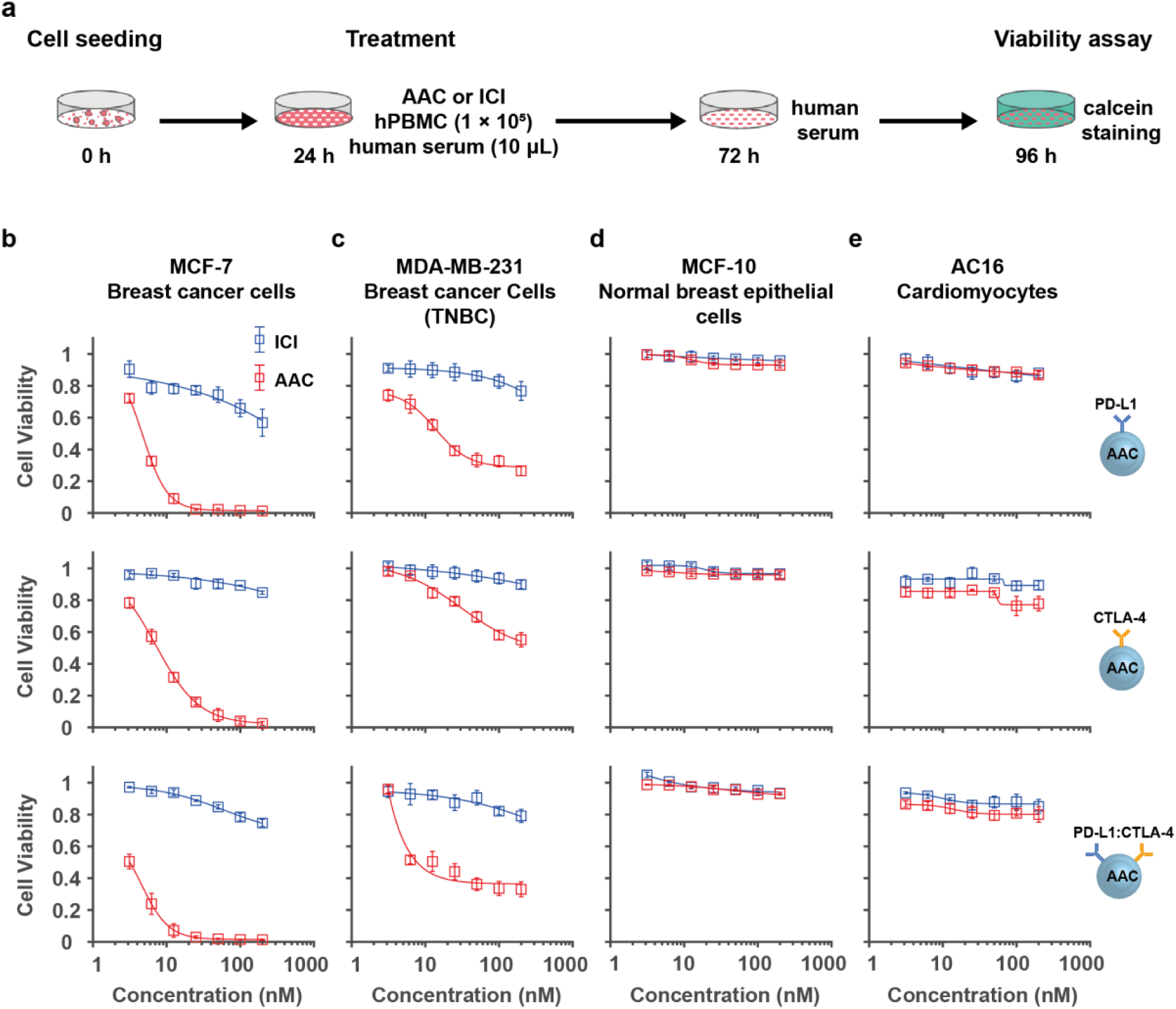
Anticancer efficacy and toxicity of AAC in cancer cells and normal cells. **a**, Schematic design of the *in vitro* assay. Two types of cancer cells and two types of normal cells were plated in 48-well plates and treated with AAC, ICI, or control reagents, followed by the addition of PBMCs to evaluate the efficacy and toxicity of the treatments in the cells. **b-e,** Viability assessments of MCF-7 (**b**) and MDA-MB-231 (**c**) cancer cells and normal MCF-10 (**d**) and AC16 (**e**) cells after treatment with anti-PD-L1 (open blue rectangle) or anti-PD-L1:AAC (open red rectangle) (top), anti-CTLA-4 (open blue rectangle) or anti-CTLA4:AAC (open red rectangle) (middle), and combo-ICI (anti-PD-L1 and anti-CTLA4) (open blue rectangle) or bi-AAC (anti-PD-L1 and anti-CTLA4) (open red rectangle) (bottom), respectively. Cell viability in each group was normalized to cell viability after PBS treatment as the basal level. Note that 800-nm nanogels were used. The experiments were repeated three (MCF-7 and AC16) or four times (MDA-MB-231 and MCF-10), and the plotted data are the mean ± standard error (SE). The unpaired t test (two-sided) was used and *p* value was not adjusted for multiple comparison (Table S1).

When the mono-AACs and bi-AAC were applied to MCF-7 cancer cells in the presence of PBMCs, the cell viability dramatically decreased in comparison to that upon treatment with the single ICIs and combination of both ICIs without the AAC (Fig. 2b-e). In particular, the half maximal inhibitory concentrations (IC_50_ values; 50% cell viability) for anti-PD-L1:AAC, anti-CTLA-4:AAC, anti-PD-L1, and anti-CTLA-4 in MCF-7 cells were estimated by fitting the measured data, providing values of 4.65 nM, 7.48 nM, 429 nM, and 7580 nM, respectively, while the IC_50_ values for the bi-AAC and combination ICI treatment were 3.17 nM and 1830 nM (Table S1), respectively. In MDA-MB-231 triple-negative breast cancer (TNBC) cells, the efficacy of the ICI:AAC-based treatments was significantly less pronounced than that in MCF-7 cells, but the IC_50_ values of anti-PD-L1:AAC, anti-CTLA-4:AAC, and the bi-AAC (15.5 nM, 419 nM, and 7.71 nM, respectively) were remarkably lower than those with the corresponding ICI treatments (1140 nM, 15300 nM, and 2410 nM, respectively) (Table S1). In contrast to the results in the cancer cells, we observed that none of the treatments was toxic to normal nontumorigenic epithelial cells (MCF-10 cell line) or human cardiomyocytes (AC16 cell line) under the same culture conditions in the presence of PBMCs. This result showed that AAC-based method substantially enhanced the anticancer efficacy of the ICIs, while keeping nontoxicity to normal cells.

### The effects of physical properties of AACs on cell viability

To determine whether the physical properties of the AAC contribute to its anticancer effects (Fig. 3), we first hypothesized that the size of Conjugel would affect the viability of MCF-7 cancer cells. To this end, differently sized Conjugels with the same composition were prepared by modifying the polymerization protocol. Thus, Conjugels with four different sizes (hydrodynamic diameters of 500 nm, 700 nm, 800 nm, and 1000 nm) were used to formulate mono-AACs (with anti-PD-L1 or anti-CTLA-4) and bi-AACs (with anti-PD-L1 and anti-CTLA-4) (Fig. 3a). When compared to the ICI alone (anti-PD-L1 and anti-CTLA-4), the larger the AACs were, the more efficient their cancer cell-killing effect was, with the 800-nm and 1000-nm AACs showing similar efficacy (Fig. 3b). Notably, the IC50 values for the 500-nm, 700-nm, 800-nm, and 1000-nm anti-PD-L1:AACs and anti-CTLA-4:AACs were 281 nM, 16.5 nM, 4.65 nM, and 6.61 nM and 648 nM, 28.1 nM, 7.48 nM, and 7.18 nM, respectively (Table S1). In addition, the same trend was observed upon bi-AAC treatment, suggesting that AAC size is important for the antitumor activity of the AAC and that 800-nm AACs would be suitable for our experiments (Fig. 3b).

**Figure 3.**
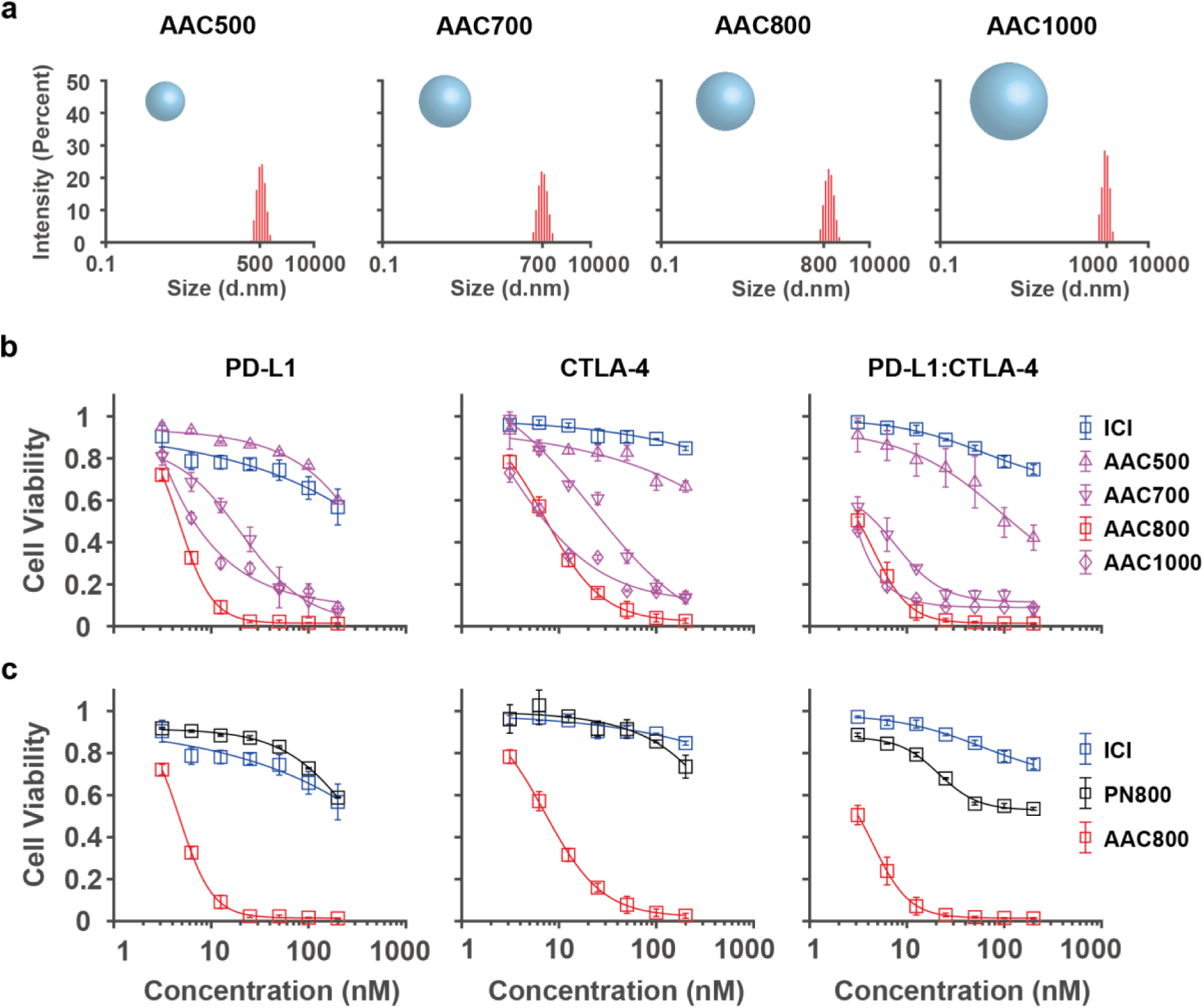
The effects of AAC size and deformability on anticancer potency. **a**, Differently sized nanogels with the same composition were prepared by modifying the polymerization protocol. The hydrodynamic diameters of the nanogels were measured by dynamic light scattering (DLS) analysis. **b,** Viability of MCF-7 breast cancer cells after treatment with single-ICI (anti-PD-L1 or anti-CTLA-4) (open blue rectangle) or various sizes of mono-AAC (anti-PD-L1 or anti-CTLA-4) (open purple triangle for 500 nm, open purple inverted triangle for 700 nm, open red rectangle for 800 nm, and open purple diamond for 1000 nm AAC) and the combo-ICI (anti-PD-L1 and anti-CTLA-4) (open blue rectangle) or various sizes of the bi-AAC (anti-PD-L1 and anti-CTLA-4) (open purple triangle for 500 nm, open purple inverted triangle for 700 nm, open red rectangle for 800 nm, and open purple diamond for 1000 nm AAC) **c,** Viability of MCF-7 breast cancer cells after treatment with single-ICI (anti-PD-L1 or anti-CTLA4) (open blue rectangle), 800-nm mono-PS (anti-PD-L1 or anti-CTLA4) (open black rectangle), or 800-nm mono-AAC (anti-PD-L1 or anti-CTLA-4) (open red rectangle) and (**d**) and the combo-ICI (anti-PD-L1 and anti-CTLA4) (open blue rectangle), 800-nm bi-PS (anti-PD-L1 and anti-CTLA4) (open black rectangle), or 800-nm bi-AAC (anti-PD-L1 and anti-CTLA4) (open red rectangle), respectively. Cell viability in each group was normalized to cell viability after PBS treatment as the basal level. The experiments were repeated three times. The plotted data are the mean ± SE. The unpaired t test (two-sided) was used for statistical analysis (Table S2).

Next, we tested whether nanogel deformability is a key factor involved in the enhanced anticancer immune response observed in the AACs vs. the ICIs alone (Fig. 3c). Using the same method previously described, nondeformable polystyrene nanoparticles (PNs) (800 nm in diameter) were conjugated with ICIs as controls for comparison to the deformable nanogel-based AACs (hydrodynamic diameter of 800 nm). The ICI alone and mono-PN (with anti-PD-L1 or anti-CTLA-4) were similarly effective in killing MCF-7 cancer cell cells (Fig. 3c), while the bi-PNs (with anti-CTLA-4 and anti-PD-L1) had a slightly improved anticancer effect (Fig. 3c). However, both the mono-and bi-AACs significantly decreased the viability of cancer cells in comparison to that upon treatment with the corresponding ICI alone and ICI-loaded PNs, respectively (Fig. 3c). Thus, we hypothesize that the size and deformability of the nanogel may regulate the AAC-mediated anticancer immune response.

### *In vivo* tests for the treatment of TNBC with AACs

Since AAC-based anticancer treatment substantially decreased the viability of cancer cells *in vitro*, we further investigated the antitumor effect of the AACs in a mouse model (Fig. 4). Mammary adenocarcinoma 4T1-luciferase (4T1-luc) cells, which resemble human TNBC cells, mixed with Matrigel were injected subcutaneously into the fourth mammary fat pad (4-MFP) of 7-week-old BALB/c female mice, which resulted in the formation of orthotopic syngeneic tumors. The antitumor efficacy of each ICI and type of AAC was estimated by measuring changes in bioluminescence intensity and tumor volume in the mice bearing 4T1-luc tumors and mice treated with saline (negative control) and nAAC (Fig. 4a-c). In single-ICI or mono-AAC treatment, anti-PD-L1 was selected because it was more efficacious than anti-CTLA-4 in the *in vitro* assay. We compared the intensity values and tumor volumes of the mono-AAC-and bi-AAC-treated mice to those of the single-ICI-and combo-ICI-treated mice every four days. On day 16 (the 12^th^ day after each treatment), mono-AAC and bi-AAC treatments had significantly different effects on bioluminescence intensity compared to those in the single-ICI and combo-ICI-treated groups (Fig. 4c and Fig. S2). The values measured from the mono-AAC-and bi-AAC-treated groups (4.3×10^5^ photons/s and 1.5×10^3^ photons/s, respectively) were ∼700-and ∼200-fold lower than those of the corresponding ICI-treated groups (3.2×10^8^ photons/s and 3.6×10^5^ photons/s), respectively, whereas both the saline (2.1×10^9^ photons/s)-and nAAC (1.7×10^9^ photons/s)-treated groups showed slightly increased intensities in contrast to those of the other groups (Fig. 4c). The changes in the volume ratios for these same groups were similar, suggesting that AAC-based ICI treatment decreased tumor burden more efficiently than treatment with the ICIs alone (Fig. S3). Moreover, the differences in bioluminescence intensity and tumor burden of these same groups were also observed at day 20 (at the point of sacrifice, see Fig. 4c and Figs. S2-S3). These results suggested that the mono-AACs and bi-AACs were superior to single-AAC and combo-ICI treatment in terms of antitumor response in 4T1-luc tumor-bearing mice. The bi-AAC was more efficacious than the mono-AAC, consistent with the *in vitro* experiments.

**Figure 4.**
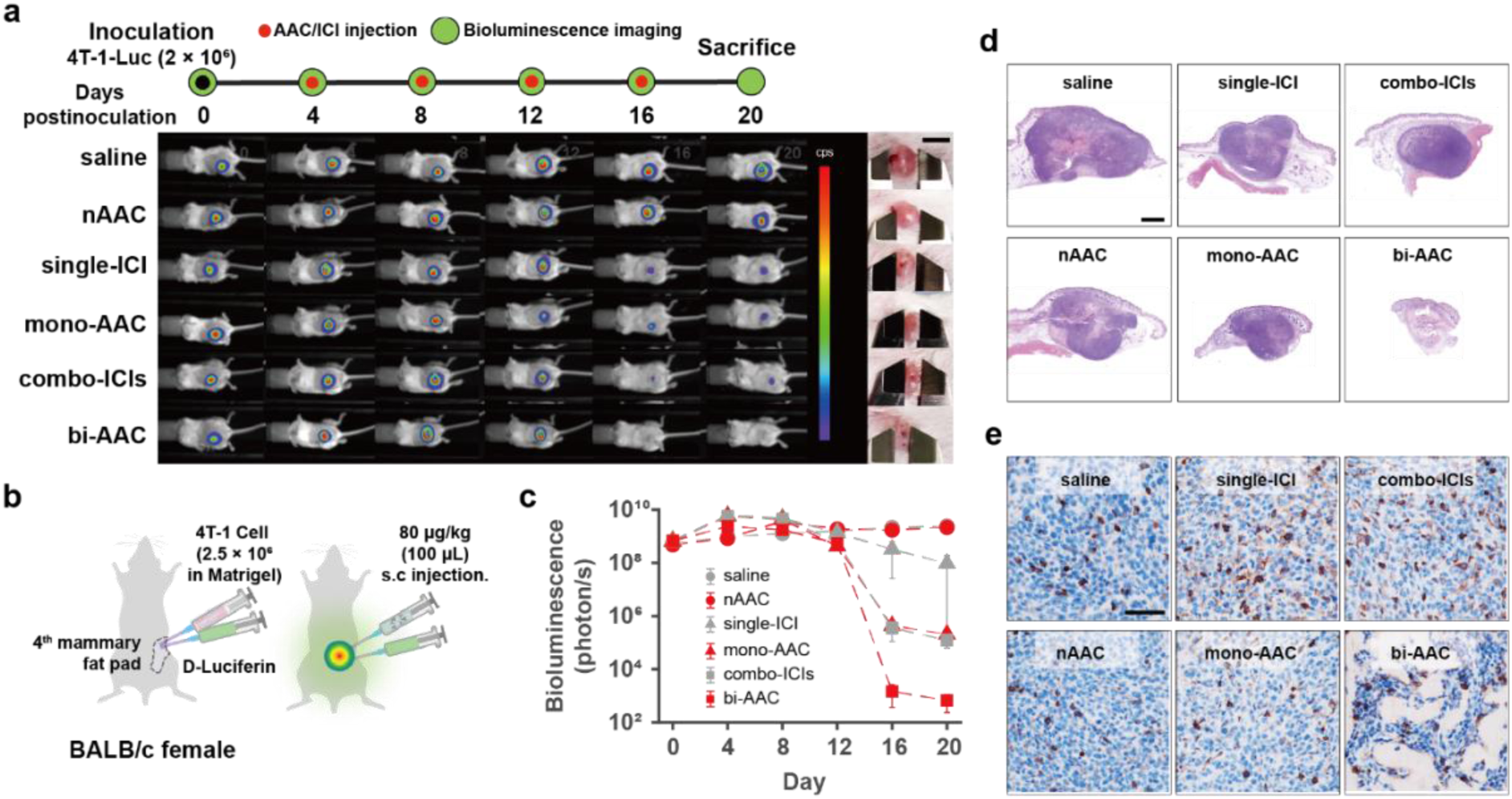
Antitumor efficacy of the ICI and AAC treatments in an orthotopic syngeneic tumor model. **a**, *In vivo* bioluminescence imaging of 4T1-luc tumor-bearing BALB/c female mice. The timeline for the experimental design (top) and representative images of the mice and tumor regions (bottom) are shown. The bioluminescence scale for each image is the same, and photographed images indicate the tumor volume on the mouse skin. The scale bar represents 5 mm. **b,** Schematic of orthotopic syngeneic tumor formation (left) and treatment (right). **c,** Bioluminescence analysis after each treatment (gray circle for saline, red circle for nAAC, gray triangle for single-ICI, red triangle for mono-AAC, gray rectangle for combo-ICI, and red rectangle for bi-AAC). The 7 mice in each group were used to estimate the mean bioluminescence signal at each time point. Error bars show the standard error (SE). The unpaired t test (two-sided) was used for statistical analysis (Table S3). **d-e,** Representative images of H&E-stained (**d**) and CD8-stained (**e**) tumor slices collected from the sacrificed mice in each treatment group. The scale bars in **d** and **e** represent 1 mm and 0.5 mm, respectively.

Next, we analyzed biopsied tumor tissues from the sacrificed mice 16 days after the treatments. Hematoxylin and eosin (H&E) staining of the sliced tumor tissues showed the morphology and cellular components of the 4T1-luc-injected area (Fig. 4d and Fig. S4). The results supported the bioluminescence observations, as the mono-AACs and bi-AACs more strongly decreased the size of the tumors than the single-ICI and combo-ICI treatment, respectively. It was also clear that bi-AAC treatment significantly reduced the tumor burden in 4T1-luc-bearing mice, since few cellular components were observed, while other treatment groups showed very dense and abundant cell nuclei, similar to control groups (Fig. 4d). Immunohistochemical analysis using an anti-CD8 antibody also confirmed that bi-AAC treatment facilitated the recruitment of CD8^+^ T cells into the tumor tissue although the statistical significance among the treated groups was not reached (Fig. 4e and Fig. S5).

To investigate whether ICI and AAC treatment altered gene expression in the tumor tissues, we performed total RNA-seq analysis of the biopsied tissues (Fig. 5). The constructed volcano plots showed that when treatment with nAAC and treatment with the bi-AAC were compared to treatment with saline as a control, the treatments altered the gene expression profile to different extents, as the expression levels of tumor microenvironment (TME)-related genes^39–43^, such as *TGFβ, TGFβR2, PLOD, CCR2, IL23,* and *COL4*, in the bi-AAC treatment group were significantly lower than those in the control group (Fig. 5a). Extended analysis of gene expression showed that in comparison to nAAC treatment, the bi-AAC treatment clearly altered expression of genes related to angiogenesis, the extracellular matrix, the immune response, and *TGFβ* compared with that in all other treated groups (single-ICI, mono-AAC, and combo-ICI) (Fig. 5b). Thus, the results of RNA-seq suggested that the effects of bi-AAC treatment were mediated via changes in the expression of a core group of genes involved in tumor-related processes.

**Figure 5.**
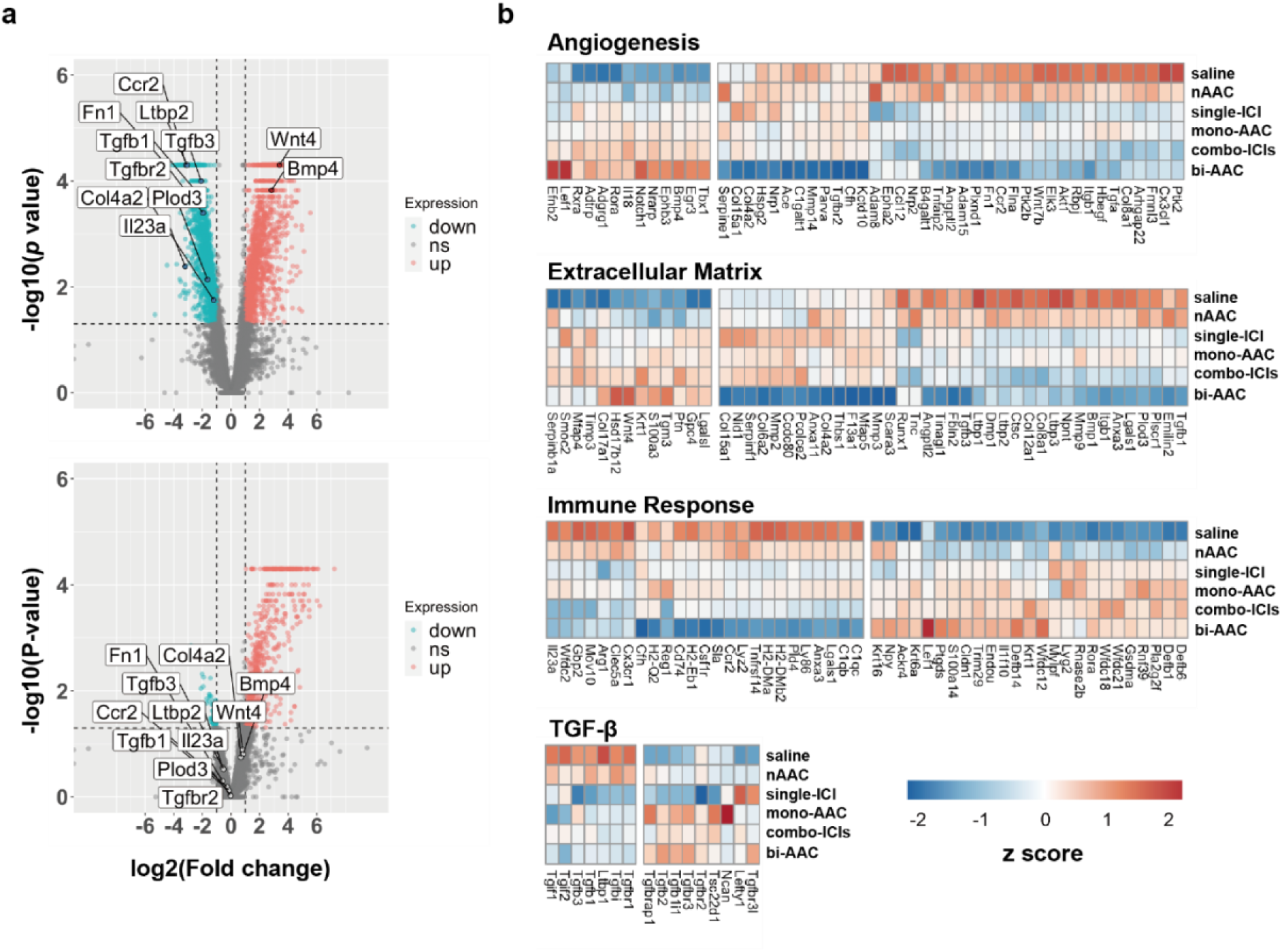
Total mRNA sequencing analysis of tumor tissue collected from the sacrificed mice after each treatment. Tumor tissue from each group (n=2) was used for mRNA sequencing analysis. **a,** Volcano plots of differentially expressed genes in the bi-AAC-and nAAC-treated groups compared to expression in the saline group are shown with annotated genes related to cancer. The dots denote upregulated genes (pastel red), downregulated genes (mint green), and genes whose expression did not significantly differ (gray). The horizontal and vertical dashed lines indicate a *p* value = 0.05 and Log_2_ FC ± 1.5. Annotated genes related to cancer were altered by bi-AAC treatment. The plot shows the – log10 *p* values and log_2_ FC values of the expressed gene mean values. **b,** The heatmaps show the representative expression of genes related to angiogenesis, the extracellular matrix (ECM), the immune response, and TGF-β in the tumor tissues collected from each treated group. Note that the heatmaps were generated by hierarchical clustering based on the Euclidean distance using the z score.

### Blood tests of mice for AAC toxicity

Finally, the biosafety of the AAC was evaluated by measuring serum concentrations of aspartate aminotransferase (AST), alanine transaminase (ALT), lactate dehydrogenase (LDH), and creatine phosphokinase (CPK) in the sacrificed mice (Fig. 6). Compared with all other groups, the bi-AAC group showed similar or lower AST, ALT, LDH, and CPK levels (Fig. 6a). For instance, AST, LDH, and CPK levels in the bi-AAC-treated mice (130 U/L, 680 U/L, and 140 U/L) were lower than those in the combo-ICI group (210 U/L, 1100 U/L, and 350 U/L, respectively). In addition, the effect of nAAC in nontumor-bearing mice was investigated by blood tests on the 20^th^ and 60^th^ days, which also showed no significant differences in these values after saline or nAAC treatment (Fig. 6b). Taken together, our results suggest that the bi-AAC may be a platform to not only maximize the therapeutic effects of ICIs but also minimize adverse effects for clinical applications.

**Figure 6.**
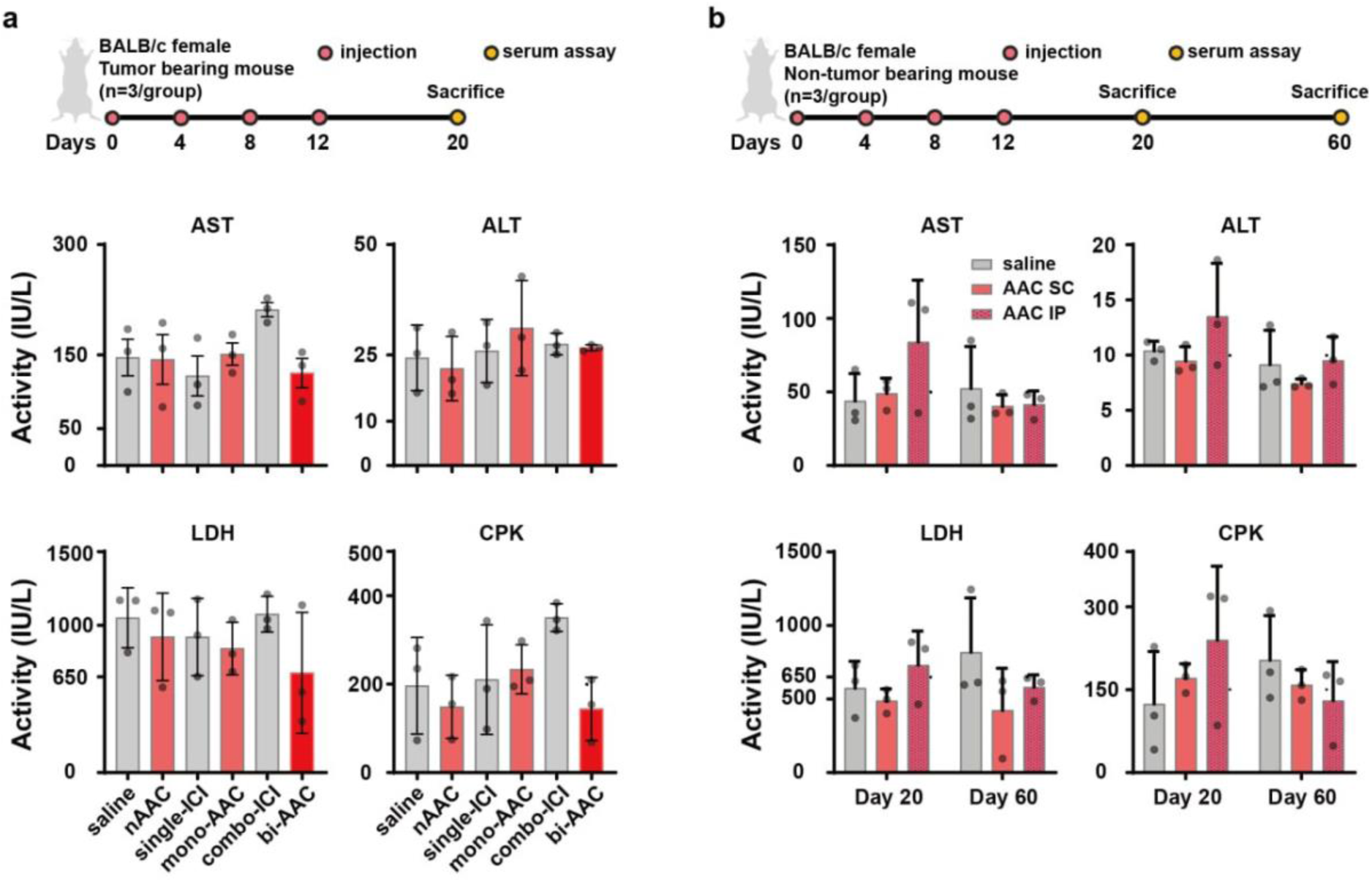
Blood tests of the AAC treated mice. **a**, Toxicity tests of blood collected from the sacrificed mice (4T1-luc tumor-bearing BALB/c female mice) in each treatment group (Fig. 5) were performed by measuring the values of AST, ALT, LDH, and CPK shown as the mean ± SD (n=3 for each group). **b,** Toxicity of nAAC was further estimated by subcutaneously (AAC SC) or intraperitoneally (AAC IP) injected into to BALB/c female mice without tumor cell inoculation (n = 3 for each group). The mice were sacrificed on day 20 or day 60 for the blood collection. The values of AST, ALT, LDH, and CPK were measured from the blood and are shown as the mean ± SD (gray bar for saline, light red bar for AAC SC, and textured red bar for AAC IP, in order).

## Discussion

We have demonstrated that compared to ICIs alone, ICIs in complex with an Conjugel had greater anticancer effects that resulted in the decreased survival of cancer cells in an *in vitro* assay and the ability to reduce tumor burden in an orthotopic syngeneic mouse model. The RNA-seq analysis also revealed that the bi-AAC treatment significantly altered the expression profile of tumor-and TME-related genes. Finally, we also showed that the AAC with ICI was not only non-toxic against the normal cells (MCF-10 and AC16) *in vitro* but also highly stable and effective for the treatment of TNBC model *in-vivo*.

Our finding that a deformable nanogel could safely enhance the efficacy of ICIs is unprecedented in cancer immunotherapy delivered via nanoparticles. Polymeric nanoparticles are typically utilized to deliver and then release drug compounds at the targeted site, but these nanoparticles only act as an efficient cargo system and do not support therapeutic synergy. Because the deformability and size of the AAC were closely related to its ability to enhance the anticancer effects of the loaded ICIs, we suggest that by modulating the biophysical conformations of nanogels, AACs can increase the therapeutic potential of ICIs for clinical treatments with improved outcomes as well as for resistant cancers.

ICI-mediated cancer immunotherapy is a ground-breaking concept in clinical oncology in which native immune cells are roused to fight cancer cells. Furthermore, CAR-T cell therapy, a strategy used to invoke such immune cell awakening by genetic engineering, has pushed immunotherapy to the next level by improving efficacy, but it comes with an extremely high medical cost. CAR-T cell therapy represents an exceptional framework for DDSs, since in this therapy, the membrane surface of living cells is embedded with therapeutic fragments or therapeutic fragments are embedded near the membrane surface. Similarly, an AAC could provide a cell-like scaffold that leverages its biophysical merits for cancer immunotherapy without the need for complex, high-cost production. Notably, nanoparticle-conjugated proteins are less susceptible to enzymatic degradation than the proteins alone, yielding a longer retention time *in vivo* than that of the soluble proteins by themselves.

In terms of avidity, antibodies immobilized on a nanoparticle or cell surface and their soluble counterparts primarily vary due to the difference in their local concentrations. When bound to epitopes on cells, surface-immobilized antibodies would be favored over soluble antibodies due to their higher avidity, which would allow them to re-engage sites at which binding does not occur. In bifunctional AACs, each antibody is thermodynamically present at the same concentration, which is an additional merit of surface immobilization that can be adjusted by altering the antibody ratio in AAC formulation. It is therefore reasonable to propose that ICIs immobilized on an AAC may have an advantage over free ICIs due to their higher avidity, which resulted in the improved anticancer effects observed in our study. Interestingly, in a clinical study of combo-ICI, a better therapeutic outcome was observed in cancer patients, but adverse effects led to the discontinuation of treatment^1^. This further justifies the potential use of AAC system in cancer immunotherapy because the blood test of experimental animals implies that the bi-AAC treatment is similar or lesser in toxicity to organs compared to the combo-ICI (Fig. 6). The results are probably due to localization of AACs at the site of tumor region through subcutaneous injection (Fig. S6) and thereby attenuation of undesired side effect by ICI in whole body. Note that unconjugated Alexa488 fluorescent dye was quickly diffused from the injected region (<1 hour), while AAC-conjugated one was stable for more than 60 days at the site (Fig. S6).

The development of bi-or multispecific antibodies that are composed of two or more paratopes is a novel concept in that artificially engineered proteins can target and regulate different kinds of biomolecules and thus their corresponding pathways simultaneously. However, to fully achieve ideal functionality, the participating moieties must be suitably spaced and able to cooperate with each epitope, qualities that are often limited by the expression and purification of an artificial protein. In contrast, the formation of bi-or multispecific antibody-like formulations in AAC systems is not limited by these requirements. It will be interesting to explore whether the AAC-based treatment can enhance the anticancer effect of ICIs in other types of cancer cells, including prostate and colon cancer cells, and whether different combinations of ICIs or mAbs can show further improved efficacy of cancer immunotherapy.

The total RNA-seq analysis of biopsied tumor tissue has been utilized to understand the changes in gene expression by cancer treatment. Our result showed that bi-AAC treatment induced apparent perturbation of gene expressions, especially related with tumorigenesis and TME. For instance, low expression of *TGFβ* and *TGFβR2* is favorable for breast cancer patient survival^44, 45^. Similarly, the expression of *CCR2, IL23*, and *COL4* is correlated with cancer progression^40–42^, and therefore, the lower expression of these genes in the bi-AAC treated group is consistent with the significantly decreased tumor burden. Furthermore, the result also implies that the adoption of other antibodies to equip the AAC may be an additional option for the follow-up treatment of relapsing breast cancers.

In conclusion, our findings suggest that the biophysical and practical merits of the AAC system can accelerate the development of many AAC-based therapeutics to more precisely treat pathological conditions, including breast cancers.

## Materials and Methods

### Animals

Female BALB/c mice (6 weeks old) were purchased from Orient Bio (Seong-nam, Korea) and were acclimated in an animal room for 1 week at 22 ± 1 °C on a 12 h light-dark cycle with 50 ± 5% humidity. The same conditions were used for all animal experiments. All animal experiments were approved by the Korea University IACUC (approval number: KOREA-2020-0203) and complied with the Code of Animal Ethics for Animal Testing and Research (Korea University, Seoul, Korea).

### Cell lines and cell culture

Two types of human breast cancer cells, MCF-7 (30022) and MDA-MB-231 (30026), were purchased from the Korean Cell Line Bank (Seoul, Korea). The mouse breast cancer cell line 4T1-Luc2 (CRL-2539-LUC2™) and the normal human epithelial cell line MCF-10A (CRL-10317™) were purchased from American Type Culture Collection (ATCC, Manassas, VA). The human cardiomyocyte cell line AC16 was purchased from Merck Millipore (Burlington, MA). Human peripheral blood mononuclear cells (PBMCs, SER-PBMC-200-F) and human serum (HSER-200ML) were purchased from Zenbio (Durham, NC).

Human MCF-7, MDA-MB-231, and AC16 cells were cultured in DMEM (including 584 mg/L L-glutamine) supplemented with 10% (v/v) fetal bovine serum and 1% (v/v) antibiotics, including streptomycin, amphotericin, and penicillin. MCF-10A cells were cultured using an MEGM™ Mammary Epithelial Cell Growth Medium BulletKit™ without gentamycin-amphotericin B mix (Lonza) following ATCC’s instructions. All cells were cultured in a humidified atmosphere at 37 °C under 5% CO_2_.

### Proteins and antibodies

Anti-human CTLA-4 (BE0190), anti-mouse CTLA-4 (BE0131) and anti-mouse PD-L1 (BE0101) antibodies were purchased from Bioxcell (Lebanon, NH). Anti-human PD-L1 antibody (10084-R639), anti-human PD-1 antibody (10377-HN94), and protein A (LC12N00802) were obtained from Sino Biological Inc. (Beijing, China). Anti-CD8 alpha antibody (ab217344) was purchased from Abcam (Cambridge, UK).

### Chemicals

All chemical compounds were used as received unless otherwise stated. Matrigel^®^ matrix was purchased from Corning (Glendale, AZ). *N*-Isopropylacrylamide (NIPAm), acrylic acid (AAc), *N,N′*-methylenebisacrylamide (BIS), ammonium persulfate (APS), poly(oxyethylene) sorbitan monooleate (Tween 80), N-hydroxysuccinimide (NHS), 1-ethyl-3-(3-dimethylaminopropyl) carbodiimide (EDC), trypan blue, and DMSO were purchased from Sigma‒Aldrich (St. Louis, MO). Alexa Fluor™ 488 hydrazide and HIER buffer H were obtained from Thermo Fisher Scientific Inc. (Waltham, MA). DMEM and PBS were purchased from Welgene (Gyeongsan, Korea) and MES was obtained from Biosolution (Seoul, Korea). Calcein AM was purchased from Invitrogen™ (Waltham, MA). Serum-free RPMI 1640 and protein block (serum-free) were obtained from Gibco (Waltham, MA) and Dako (Santa Clara, CA), respectively.

### Synthesis of pNIPAm-AAc nanogels and dynamic light scattering (DLS) analysis

Deformable nanogels were prepared using aqueous free-radical precipitation polymerization as previously described^46, 47^. Briefly, an 88% pNIPAM/10% AAc/2% BIS solution (poly(*N*-isopropylacrylamide-*co*-acrylic acid)) was used for nanogel synthesis. The reaction mixture was prepared by dissolving NIPAm and BIS in deionized water, followed by the addition of Tween 80 or SDS as a surfactant. The mixture, contained in a reaction vessel, was heated at 70 °C and purged with nitrogen gas. After 1 h, a solution of AAc was added to the mixture, followed by 10 min of nitrogen gas purging. The reaction was initiated by adding APS solution to the vessel and carried out for 6 h at 70 °C under nitrogen gas purging. At the end of the reaction, the polymerized nanogels were purified by dialysis against deionized water for 1 week by changing the water twice per day. The purified nanogels were lyophilized before storage.

### AAC preparation

Nanogels (10 mg/mL) resuspended in 0.1 M MES buffer (pH=5.5) were mixed with EDC (20 mg/mL) and NHS (40 mg/mL) to a final nanogel concentration of 5 mg/mL. After 5 min of preincubation, protein A was added to the mixture, at a final concentration of 10 µM, and the mixture was incubated for 1 h at RT. After protein A coupling via EDC/NHS, the mixture was centrifuged for 2 min at 8000 rpm using a table top centrifuge, and the supernatant was exchanged with PBS buffer. This procedure was performed 5 times to remove unreacted protein A, EDC, and NHS. The protein A-functionalized nanogel (nAAC), we call this the “Conjugel”. was stored at 4 °C until the *in vitro* and *in vivo* experiments were performed. To prepare mono-AACs, nAAC (5 mg/mL) was mixed with an equivalent volume of anti-PD-L1 antibody (8 µM) or anti-CTLA-4 antibody (8 µM), while the bi-AAC was prepared by mixing nAAC (5 mg/mL) with anti-PD-L1 (8 µM) and anti-CTLA-4 (8 µM) antibodies at a 2:1:1 volume ratio.

To prepare the fluorescently labeled AACs (Fl-AACs), nanogel (10 mg/mL) resuspended in 0.1 M MES buffer (pH=5.5) was mixed with an amine-reactive fluorescent dye (Alexa Fluor™ 488 hydrazide) (2 mM in DMSO) and EDC (100 mg/mL), followed by overnight incubation at RT. The mixture was centrifuged for 5 min at 13,000 rpm using a table top centrifuge, and the supernatant was exchanged with 0.1 M MES buffer (pH=5.5). This procedure was performed 5 times to remove unreacted dye, MES, and DMSO. After fluorescent labeling of the nanogel, protein A coupling was performed as described above, and then the Fl-AAC was incubated with 400 nM anti-PD-L1 antibody or isotype antibody.

### AAC treatment for cell viability assays

MCF-7 (1 × 10^4^ cells/well), MDA-MB-231 (1 × 10^4^ cells/well), MCF-10A (1 × 10^3^ cells/well), and AC16 (5 × 10^3^ cells/well) cells were plated in 48-well plates and incubated for 24 h before treatment. On day 0, negative control reagents, ICIs, or AACs diluted in DMEM (100 µL) were added to each well, followed by the addition of 1 × 10^5^ hPBMCs (resuspended in 100 µL of DMEM) and 10 µL of human serum. At 72 h, 20 µL of DMEM and 10 µL of human serum were added to each well to compensate for evaporation of the culture medium. At 96 h, the culture medium was removed, and cell debris was washed away twice with prewarmed PBS. The remaining cells were stained by treatment with 200 µL of the fluorescent live-cell dye calcein AM (0.5 µg/mL), followed by incubation at 37 °C for 15 min avoiding light. The dye was removed from each well and the cells were washed twice with prewarmed PBS, after which the wells were filled with medium for imaging analysis. Fluorescence microscopy images were captured by a Nikon Eclipse Ti2 (Nikon Instrument Inc. NY) with a 4× objective, 10× eyepiece, and excitation filter (EX465-495). Cell viability was estimated by analyzing the fluorescent region of the captured images with ImageJ software.

### Cell localization via assessment of Fl-AACs

MDA-MB-231 cells were plated in an 8-well chamber (1 × 10^4^ cells/well) (Nunc™ Lab-Tek™ II chambered cover glass) and cultured for three days. Then, the culture medium was removed, and the cells were washed twice with prewarmed PBS to remove cell debris. Cells were fixed in 3% paraformaldehyde (PFA) with 0.1% glutaraldehyde diluted in PBS for 30 min. The cell membranes were stained with lipophilic CM-DiI dye (5 µg/mL, Vybrant™ CM-DiI Cell-Labeling Solution, Invitrogen) at RT for 15 min.

Fl-AAC-PD-L1 or Fl-AAC-isotype was added to the cells (200 µL/well), followed by incubation overnight at 4 °C. The cells were washed twice with PBS and stained with DAPI (0.5 µg/mL) (Roche) for 10 min at RT. Fluorescence microscopy images were captured by a Nikon Eclipse Ti2 with a 40× objective and the corresponding excitation filters. The fluorescence signal density (area × intensity) of the captured images was analyzed with ImageJ software to estimate the effective targeting of AAC-PD-L1 to cancer cells.

### Animal experiments

Six-week-old BALB/c female mice (n=7 per group) were acclimated in an animal room for one week before treatment. The experimental groups were defined as follows based on the materials that were administered: (1) saline, (2) nAAC, (3) single-ICI, (4) mono-AAC, (5) combo-ICI and (6) bi-AAC.

For orthotopic syngeneic tumor modeling, the mice (at 7 weeks of age, 15∼18 g) were anesthetized using 1.5% isoflurane (Hana Pharm, Co., Ltd., Seoul, Korea) inhaled through a mask. 4T1-luc cells (2.5 × 10^6^) resuspended in 150 µL of serum-free RPMI 1640 (Gibco) were mixed with 50 µL of Matrigel^®^ matrix (18.33 mg/mL), and the mixture was inoculated into the 4^th^ mammary fat pad of the mice (Day 0). On days 4, 8, 12 and 16 after inoculation (Day 0), the mice were treated with (1) 100 *µ*L of 0.9% normal saline, (2) 15.54 µL of nAAC, (3) 8.36 µL of anti-PD-L1 mAb (3) in saline (single-ICI), (4) 15.54 µL of AAC containing anti-PD-L1 mAb (3 mg/kg), (5) 8.36 µL of anti-PD-L1 mAb (3 mg/kg) and 7.18 µL of anti-CTLA-4 mAb (3 mg/kg) as a mAb mixture (combo-ICI), or (6) 15.54 µL of nAAC containing anti-PD-L1 mAb (3 mg/kg) and anti-CTLA-4 mAb (3 mg/kg) (bi-AAC). The injection volume was 100 µL after mixing with saline. Tumor volume (1/2 × longitudinal diameter × transverse diameter^2^) was calculated based on measurements with a digital Vernier caliper at each treatment time.

### *In vivo* bioluminescence imaging

For bioluminescence analysis of the tumor burden, the treated mice were subcutaneously injected with 100 µL of D-luciferin, potassium salt (80 µg/kg) (Goldbio, MO) 15 min before each observation. To determine tumor burden, the luciferase activity was detected using NightOWL LB 981 (Berthold Technologies GmbH & Co. KG, Bad Wildbad, Germany) when the mice were anesthetized with continuous delivery of 4% isoflurane via a nose cone system. *In vivo* imaging was performed every four days for 20 days. For fluorescence time-course analysis of AAC or free dye, Fl-AAC or Alexa488 dye was subcutaneously injected in saline and *in vivo* imaging was performed every four days for 60 days or 20 days over an hour. The captured images were analyzed using IndiGO software 2.0.5.

### Measurements of serum AST, ALT, LDH and CK

Blood (500∼700 µL) was collected from the left cardiac atrium of each mouse and then centrifuged at 3000 rpm for 15 min to separate the serum. Serum levels of aspartate aminotransferase (AST) and alanine aminotransferase (ALT) were measured using the Asan Set GOT and GPT Assay Kit (Asan Pharmaceuticals Co., Ltd., Seoul, Korea), and lactate dehydrogenase (LDH) and creatine kinase (CK) levels were assessed using the Lactate Dehydrogenase Activity Assay Kit and Creatine Kinase Activity Assay Kit (Sigma), respectively. Quantification of the results was carried out with a multiplate reader (Spectramax iD3, Molecular Devices, CA).

### Tissue harvesting

On the 20^th^ day, the mice were sacrificed by CO_2_ administration. Tumor tissues (n = 7) were harvested, and two of them were bisected and fixed in 4% PFA for histopathological analysis. The other tissue samples were preserved by snap freezing using liquid nitrogen, and two samples were used to perform total mRNA sequencing analysis.

### Histopathological analysis

Immunohistochemical and hematoxylin and eosin (H&E) staining was performed on sliced sections (3 *µ*m) that had been formalin-fixed/paraffin-embedded (FFPE) using an autostainer (Thermo Fisher Scientific Inc.). For immunohistochemistry (IHC), deparaffinization and antigen retrieval were carried out with the PT module using dewax and HIER buffer H, followed by the addition of serum-free protein block to inhibit nonspecific staining. Tissue samples were incubated for 3 h at RT with the primary antibody (anti-CD8 alpha) before visualization with the Polink-2 HRP Plus Broad DAB Detection System for Immunohistochemistry kit (GBI Labs, Bothell, WA) according to the manufacturer’s instructions. H&E-stained sections were imaged by a Nikon Eclipse Ti2 (Nikon Instrument Inc. NY) equipped with a color camera (HK20E3S, KOPTIC, Yongin, Korea). The CD8-positive ratio was assessed by dividing the cell area by the DAB-positive area in each of the images using ImageJ software.

## Total mRNA sequencing of tumor tissue

### RNA isolation

Total RNA was isolated from the tumor tissues using TRIzol reagent (Invitrogen). RNA quality was assessed by a TapeStation4000 system (Agilent Technologies, Amstelveen, Netherlands), and RNA quantification was performed using an ND-2000 spectrophotometer (Thermo Inc., DE, USA).

### Library preparation and sequencing

Libraries were prepared from total RNA using the NEBNext Ultra II Directional RNA-Seq Kit (New England BioLabs, Inc., UK). mRNA isolation was performed using the Poly(A) RNA Selection Kit (LEXOGEN, Inc., Austria). The isolated mRNA was used for cDNA synthesis and shearing following the manufacturer’s instructions. Indexing was performed using Illumina indexes 1-12. Enrichment was carried out using PCR. Subsequently, the libraries were checked using TapeStation HS D1000 Screen Tape (Agilent Technologies, Amstelveen, The Netherlands) to evaluate the mean fragment size. Quantification was performed using a library quantification kit and a StepOne real-time PCR system (Life Technologies, Inc., USA). High-throughput sequencing was performed as paired-end 100 sequencing using NovaSeq 6000 (Illumina, Inc., USA).

### Sequencing data analysis

Quality control of the raw sequencing data was performed using FastQC (Simon, 2010). Reads containing adapter sequences and low-quality reads (< Q20) were removed using FASTX_Trimmer (Hannon Lab, 2014) and BBMap (Bushnell, 2014). Then, the trimmed reads were mapped to the reference genome using TopHat (Cole Trapnell et al., 2009). The read count (RC) data were processed based on the FPKM+geometric normalization method using EdgeR within R (R development Core Team, 2020). Fragments per kb per million reads (FPKM) values were estimated using Cufflinks (Roberts et al., 2011). Data mining and graphic visualization were performed using ExDEGA (Ebiogen Inc., Korea).

### Statistical analysis

Data are presented as the mean ± standard deviation (SD) or standard error (SE). Significance was defined based on *p* < 0.05 as assessed by unpaired t-test. Each figure legend includes the detailed corresponding statistical methods. All statistical data were calculated using Matlab (Mathworks, Natick, MA, USA).

## References

1. Wolchok, J.D. et al. Overall Survival with Combined Nivolumab and Ipilimumab in Advanced Melanoma. N Engl J Med 377, 1345–1356 (2017).

2. Chae, Y.K. et al. Current landscape and future of dual anti-CTLA4 and PD-1/PD-L1 blockade immunotherapy in cancer; lessons learned from clinical trials with melanoma and non-small cell lung cancer (NSCLC). J Immunother Cancer 6, 39 (2018).

3. Sharma, P. & Allison, J.P. Immune checkpoint targeting in cancer therapy: toward combination strategies with curative potential. Cell 161, 205–214 (2015).

4. Hodi, F.S. et al. Combined nivolumab and ipilimumab versus ipilimumab alone in patients with advanced melanoma: 2-year overall survival outcomes in a multicentre, randomised, controlled, phase 2 trial. Lancet Oncol 17, 1558–1568 (2016).

5. Gao, J. et al. Neoadjuvant PD-L1 plus CTLA-4 blockade in patients with cisplatin-ineligible operable high-risk urothelial carcinoma. Nat Med 26, 1845–1851 (2020).

6. Veatch, J.R. & Riddell, S.R. Immune checkpoint blockade provokes resident memory T cells to eliminate head and neck cancer. Cell 185, 2848–2849 (2022).

7. Marin-Acevedo, J.A., Kimbrough, E.O. & Lou, Y. Next generation of immune checkpoint inhibitors and beyond. J Hematol Oncol 14, 45 (2021).

8. Ludin, A. & Zon, L.I. Cancer immunotherapy: The dark side of PD-1 receptor inhibition. Nature 552, 41–42 (2017).

9. Morad, G., Helmink, B.A., Sharma, P. & Wargo, J.A. Hallmarks of response, resistance, and toxicity to immune checkpoint blockade. Cell 184, 5309–5337 (2021).

10. Wang, H. & Mooney, D.J. Biomaterial-assisted targeted modulation of immune cells in cancer treatment. Nat Mater 17, 761–772 (2018).

11. Demaria, O. et al. Harnessing innate immunity in cancer therapy. Nature 574, 45–56 (2019).

12. Larkin, J. et al. Five-Year Survival with Combined Nivolumab and Ipilimumab in Advanced Melanoma. N Engl J Med 381, 1535–1546 (2019).

13. Chau, C.H., Steeg, P.S. & Figg, W.D. Antibody-drug conjugates for cancer. Lancet 394, 793–804 (2019).

14. Minn, A.J. & Wherry, E.J. Combination Cancer Therapies with Immune Checkpoint Blockade: Convergence on Interferon Signaling. Cell 165, 272–275 (2016).

15. Heinhuis, K.M. et al. Enhancing antitumor response by combining immune checkpoint inhibitors with chemotherapy in solid tumors. Ann Oncol 30, 219–235 (2019).

16. Chocarro, L. et al. Cutting-Edge: Preclinical and Clinical Development of the First Approved Lag-3 Inhibitor. Cells 11 (2022).

17. Drago, J.Z., Modi, S. & Chandarlapaty, S. Unlocking the potential of antibody-drug conjugates for cancer therapy. Nat Rev Clin Oncol 18, 327–344 (2021).

18. Hirsch, I., Goldstein, D.A., Tannock, I.F., Butler, M.O. & Gilbert, D.C. Optimizing the dose and schedule of immune checkpoint inhibitors in cancer to allow global access. Nat Med 28, 2236–2237 (2022).

19. Zhang, H. et al. Regulatory mechanisms of immune checkpoints PD-L1 and CTLA-4 in cancer. J Exp Clin Cancer Res 40, 184 (2021).

20. Kjeldsen, J.W. et al. A phase 1/2 trial of an immune-modulatory vaccine against IDO/PD-L1 in combination with nivolumab in metastatic melanoma. Nat Med 27, 2212–2223 (2021).

21. Dougan, M., Luoma, A.M., Dougan, S.K. & Wucherpfennig, K.W. Understanding and treating the inflammatory adverse events of cancer immunotherapy. Cell 184, 1575–1588 (2021).

22. Martins, F. et al. Adverse effects of immune-checkpoint inhibitors: epidemiology, management and surveillance. Nat Rev Clin Oncol 16, 563–580 (2019).

23. Stadler, C.R. et al. Elimination of large tumors in mice by mRNA-encoded bispecific antibodies. Nat Med 23, 815–817 (2017).

24. Sharkey, R.M. et al. Signal amplification in molecular imaging by pretargeting a multivalent, bispecific antibody. Nat Med 11, 1250–1255 (2005).

25. Herpers, B. et al. Functional patient-derived organoid screenings identify MCLA-158 as a therapeutic EGFR x LGR5 bispecific antibody with efficacy in epithelial tumors. Nat Cancer 3, 418–436 (2022).

26. Liu, L. et al. Rejuvenation of tumour-specific T cells through bispecific antibodies targeting PD-L1 on dendritic cells. Nat Biomed Eng 5, 1261–1273 (2021).

27. Dovedi, S.J. et al. Design and Efficacy of a Monovalent Bispecific PD-1/CTLA4 Antibody That Enhances CTLA4 Blockade on PD-1(+) Activated T Cells. Cancer Discov 11, 1100–1117 (2021).

28. Riley, R.S., June, C.H., Langer, R. & Mitchell, M.J. Delivery technologies for cancer immunotherapy. Nat Rev Drug Discov 18, 175–196 (2019).

29. Goldberg, M.S. Improving cancer immunotherapy through nanotechnology. Nat Rev Cancer 19, 587–602 (2019).

30. de Lazaro, I. & Mooney, D.J. Obstacles and opportunities in a forward vision for cancer nanomedicine. Nat Mater 20, 1469–1479 (2021).

31. Haber, T. et al. Specific targeting of ovarian tumor-associated macrophages by large, anionic nanoparticles. Proc Natl Acad Sci U S A 117, 19737–19745 (2020).

32. Vargason, A.M., Anselmo, A.C. & Mitragotri, S. The evolution of commercial drug delivery technologies. Nat Biomed Eng 5, 951–967 (2021).

33. Rodell, C.B. et al. TLR7/8-agonist-loaded nanoparticles promote the polarization of tumour-associated macrophages to enhance cancer immunotherapy. Nat Biomed Eng 2, 578–588 (2018).

34. Boehnke, N. et al. Massively parallel pooled screening reveals genomic determinants of nanoparticle delivery. Science 377, eabm5551 (2022).

35. Lim, W.A. & June, C.H. The Principles of Engineering Immune Cells to Treat Cancer. Cell 168, 724–740 (2017).

36. Leach, D.R., Krummel, M.F. & Allison, J.P. Enhancement of antitumor immunity by CTLA-4 blockade. Science 271, 1734–1736 (1996).

37. Wang, T.W. et al. Blocking PD-L1-PD-1 improves senescence surveillance and ageing phenotypes. Nature 611, 358–364 (2022).

38. Yang, H.M. et al. Label-Free Analysis of Multivalent Protein Binding Using Bioresponsive Nanogels and Surface Plasmon Resonance (SPR). ACS Appl Mater Interfaces 12, 5413–5419 (2020).

39. Massague, J. TGFbeta in Cancer. Cell 134, 215–230 (2008).

40. Xu, S. et al. The role of collagen in cancer: from bench to bedside. J Transl Med 17, 309 (2019).

41. Langowski, J.L. et al. IL-23 promotes tumour incidence and growth. Nature 442, 461–465 (2006).

42. Hao, Q., Vadgama, J.V. & Wang, P. CCL2/CCR2 signaling in cancer pathogenesis. Cell Commun Signal 18, 82 (2020).

43. Winkler, J., Abisoye-Ogunniyan, A., Metcalf, K.J. & Werb, Z. Concepts of extracellular matrix remodelling in tumour progression and metastasis. Nat Commun 11, 5120 (2020).

44. Buck, M.B., Fritz, P., Dippon, J., Zugmaier, G. & Knabbe, C. Prognostic significance of transforming growth factor beta receptor II in estrogen receptor-negative breast cancer patients. Clin Cancer Res 10, 491–498 (2004).

45. Aleckovic, M. et al. Breast cancer prevention by short-term inhibition of TGFbeta signaling. Nat Commun 13, 7558 (2022).

46. Lee, S.K. et al. Method for the Rapid Detection of SARS-CoV-2-Neutralizing Antibodies Using a Nanogel-Based Surface Plasmon Resonance Biosensor. ACS Appl Polym Mater 5, 2195–2202 (2023).

47. Teoh, J.Y. et al. Tuning Surface Plasmon Resonance Responses through Size and Crosslinking Control of Multivalent Protein Binding-Capable Nanoscale Hydrogels. ACS Biomater Sci Eng 8, 2878–2889 (2022).

