## supplemental information for "Antibody-conjugating nanogel (Conjugel) with two immune checkpoint inhibitors for enhanced cancer immunotherapy"

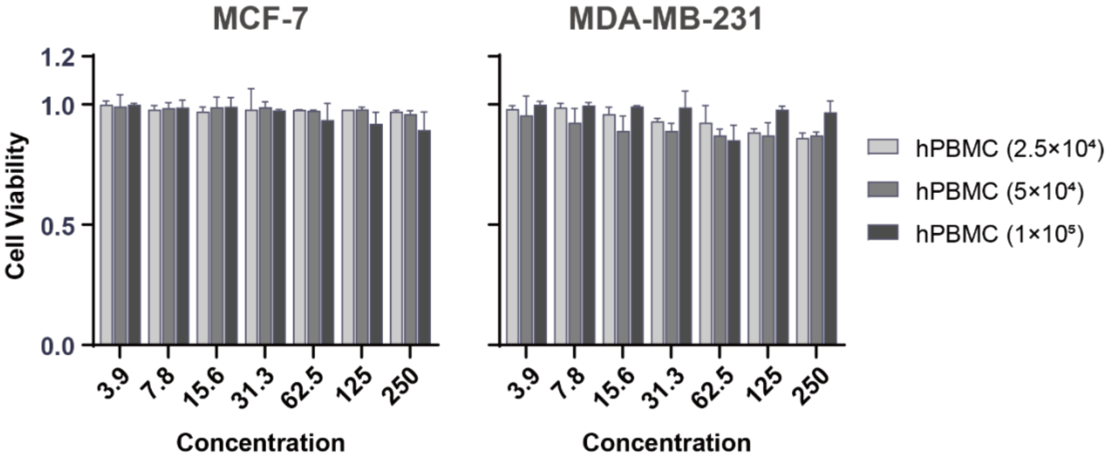


**Figure S1.** Anticancer efficacy of PBMCs and AAC without an ICI (nAAC) in two types of breast cancer cells. Note that 800 nm nanogel was used. The experiments were repeated four times and plotted by the mean ± standard deviation (s.d.).

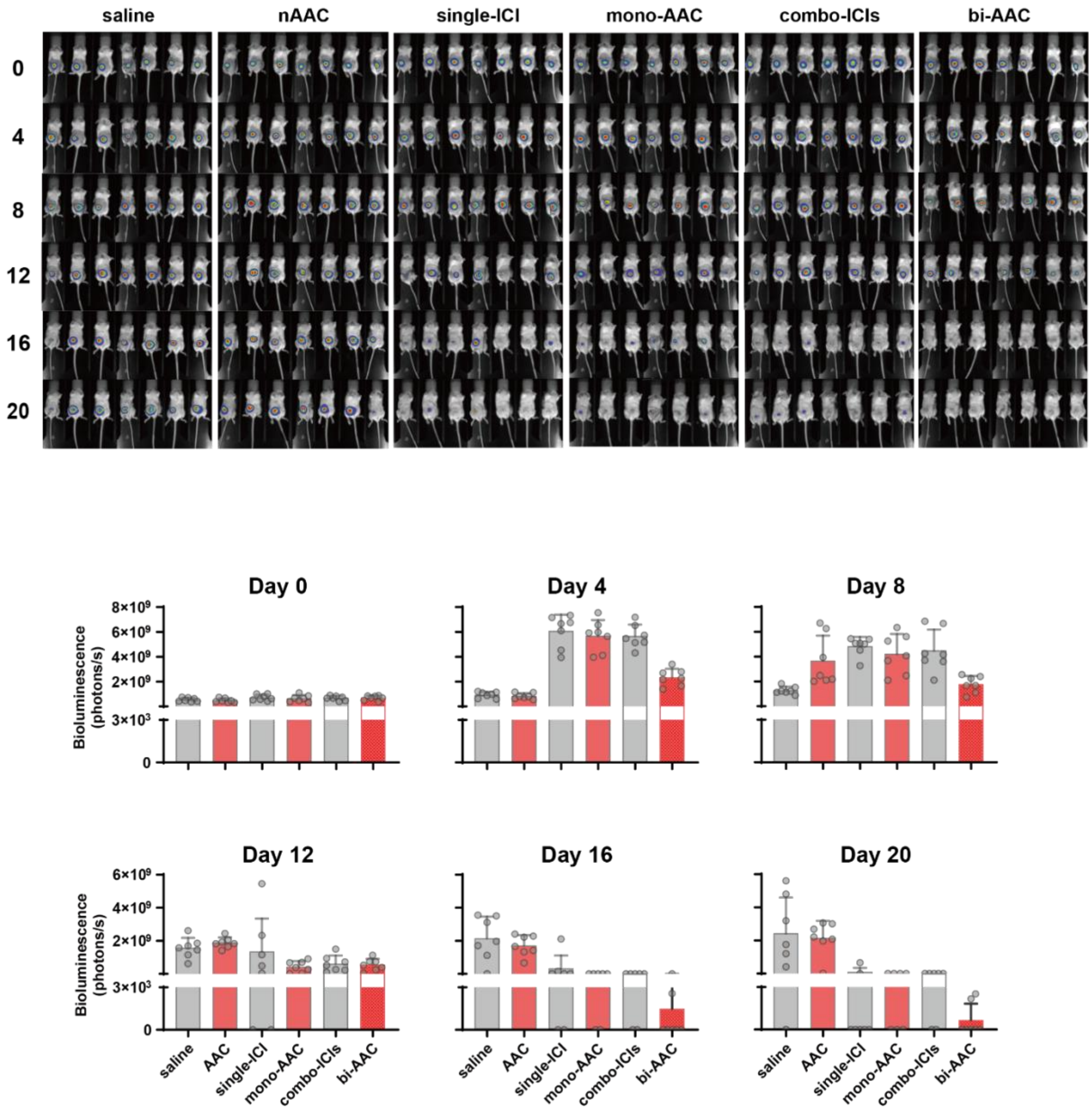


**Figure S2.** Bioluminescence intensity (images and bar graphs) of 4T1-luc tumor-bearing mice after each treatment presented in Figure 4c. Note that 800 nm nanogel was used. Error bars show the standard deviation (s.d.). Unpaired t-test was used for statistical analysis (Table S4).

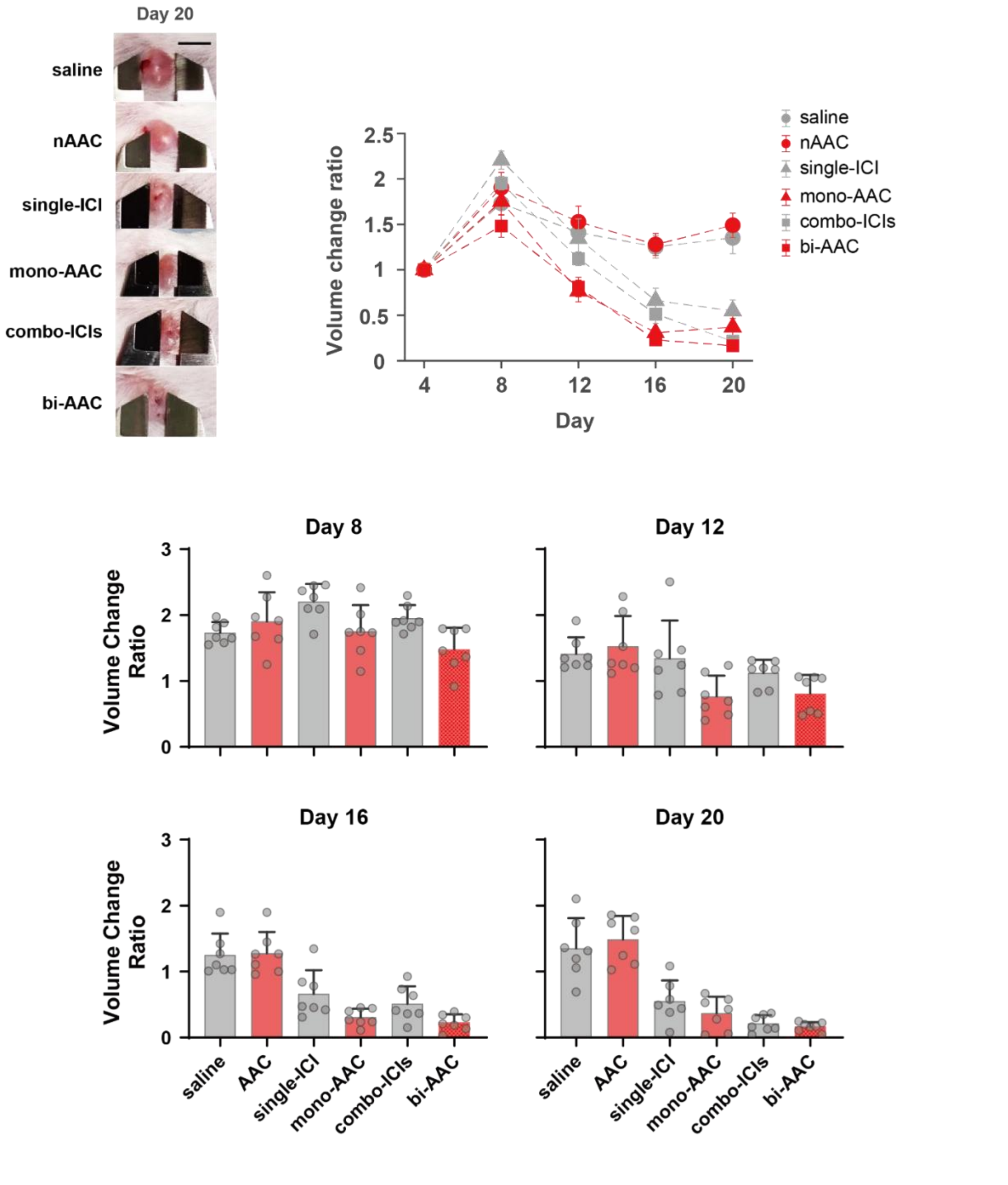


**Figure S3.** Changes in 4T1-luc tumor-bearing mouse tumor volumes after each treatment. Error bars show the standard error for the graphs and standard deviation for the histograms.

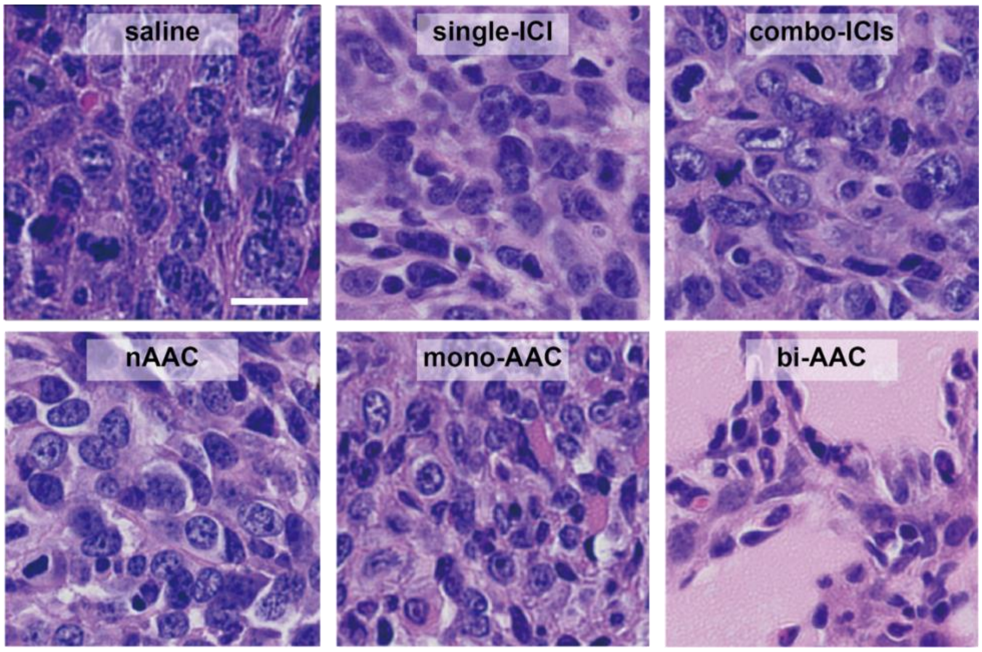


**Figure S4.** Representative zoomed images of H&E-stained tumor slices collected from the sacrificed mice in each treatment group (Fig. 4d) are presented. The scale bar is 500 μm.

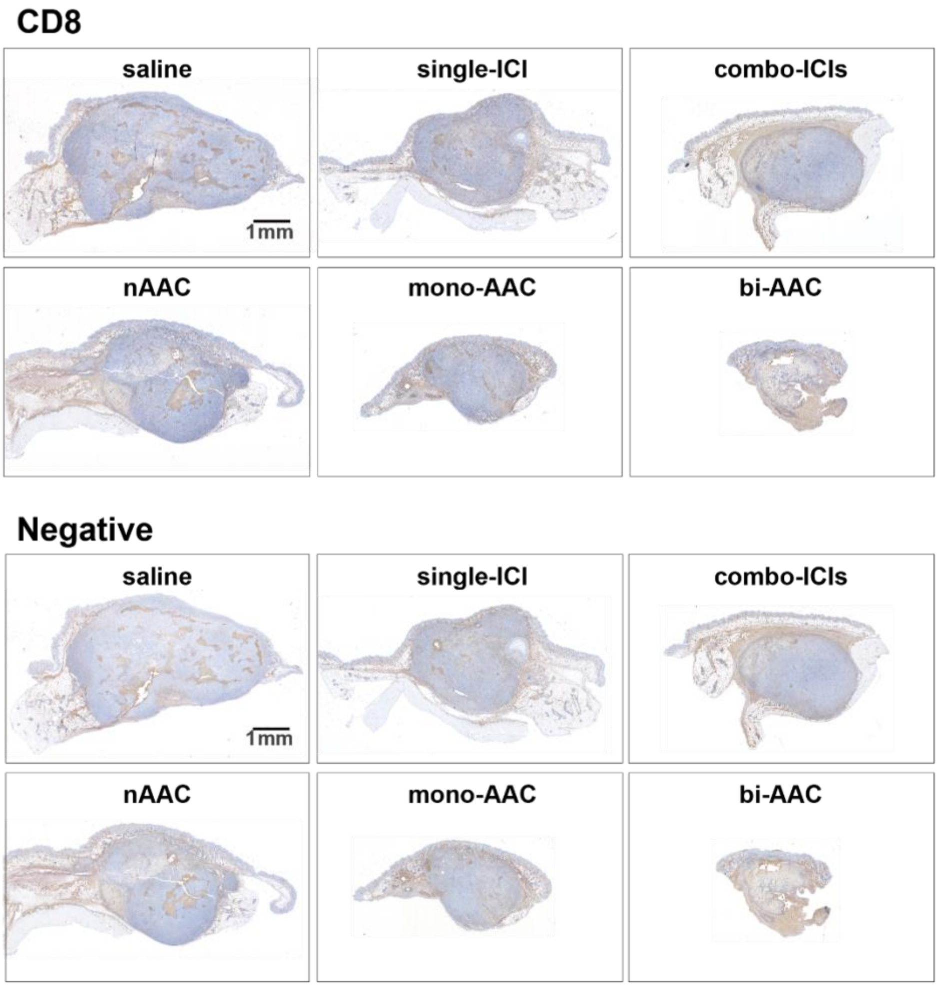


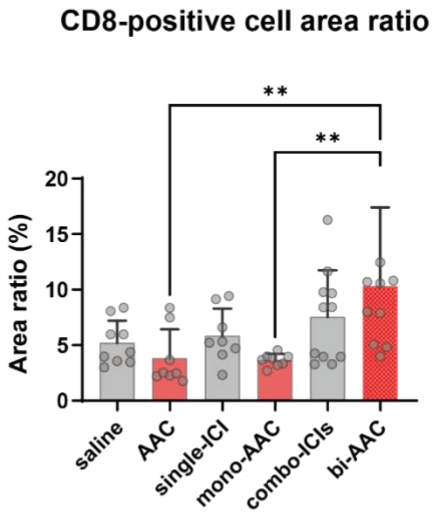


**Figure S5.** Images of CD8-stained tumor slices collected from the sacrificed mice in each treatment group. The scale bar is 1 mm. CD8-positive signal ratio was assessed by dividing the cell area by the DAB-positive area of the images. Negative control is also presented. Error bars show the standard deviation (s.d.).

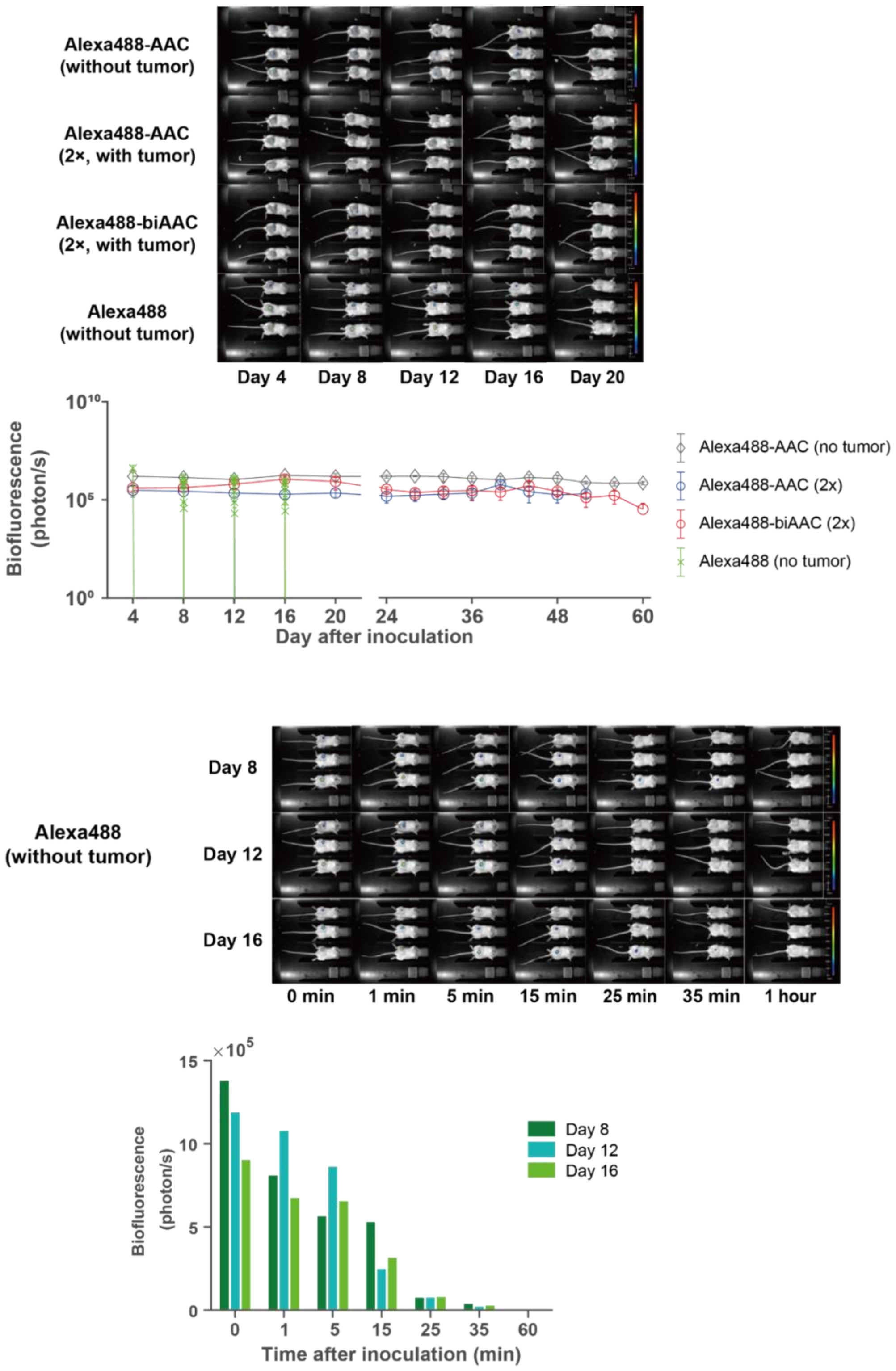


**Figure S6.** In-vivo time-course analysis of biofluorescence of AAC conjugated- or free-dye. Fl-AAC or free dye was injected into mice with different experimental conditions (n=3). The graphs on the plot above shows the long-term change in fluorescence level for each condition. The graphs on the plot below shows the short-term decay of fluorescence level right after alexa488 dye dose each day.

| AAC | MCF-7 | | MDA-MB-231 | |
|---|---|---|---|---|
| | (+) Gel | (-) Gel | (+) Gel | (-) Gel |
| PD-L1 | 4.65 | 360 | 15.5 | 870 |
| CTLA-4 | 7.48 | 5190 | 419 | 15300 |
| PD-L1:CTLA-4 | 3.17 | 1890 | 7.71 | 4300 |

| | PD-L1 | CTLA-4 | PD-L1:CTLA-4 |
|---|---|---|---|
| ICI | 524 | 5240 | 2130 |
| AAC500 | 281 | 648 | 118 |
| AAC700 | 16.5 | 28.1 | 4.69 |
| AAC800 | 4.65 | 7.48 | 3.17 |
| AAC1000 | 6.61 | 7.18 | 2.93 |
| PS800 | 282 | 456 | 207 |

**Table S1.** $IC_{50}$ values of breast cancer cells. The table above shows the difference between the values of AAC-treated cell groups and ICI-treated cell groups. The table below shows $IC_{50}$ values of different MCF-7 cell groups.

MCF-7

| Concentration (nM) | PD-L1 | CTLA-4 | PD-L1:CTLA-4 |
|---|---|---|---|
| 3.125 | 0.033589 | 0.010784 | 0.000541 |
| 6.25 | 0.000708 | 0.001128 | 0.000472 |
| 12.5 | 0.000107 | 0.000043 | 0.000047 |
| 25 | 0.000008 | 0.000055 | <0.000001 |
| 50 | 0.000125 | 0.000069 | <0.000001 |
| 100 | 0.000280 | 0.000001 | 0.000006 |
| 200 | 0.002886 | <0.000001 | 0.000006 |

MDA-MB-231

| Concentration (nM) | PD-L1 | CTLA-4 | PD-L1:CTLA-4 |
|---|---|---|---|
| 3.125 | 0.006841 | 0.306306 | 0.755042 |
| 6.25 | 0.028769 | 0.343755 | 0.001095 |
| 12.5 | 0.001073 | 0.050507 | 0.000840 |
| 25 | 0.000178 | 0.009004 | 0.000886 |
| 50 | 0.000046 | 0.002432 | 0.000100 |
| 100 | 0.000110 | 0.000225 | 0.000063 |
| 200 | 0.000271 | 0.000547 | 0.000306 |

MCF-10

| Concentration (nM) | PD-L1 | CTLA-4 | PD-L1:CTLA-4 |
|---|---|---|---|
| 3.125 | 0.967823 | 0.172315 | 0.012007 |
| 6.25 | 0.819398 | 0.111354 | 0.036914 |
| 12.5 | 0.649055 | 0.101027 | 0.822725 |
| 25 | 0.078341 | 0.523586 | 0.559250 |
| 50 | 0.038698 | 0.543792 | 0.807324 |
| 100 | 0.139102 | 0.716108 | 0.136026 |
| 200 | 0.224389 | 0.706006 | 0.784198 |

AC16

| Concentration (nM) | PD-L1 | CTLA-4 | PD-L1:CTLA-4 |
|---|---|---|---|
| 3.125 | 0.513697 | 0.250481 | 0.022513 |
| 6.25 | 0.626093 | 0.058379 | 0.094420 |
| 12.5 | 0.979193 | 0.191303 | 0.164297 |
| 25 | 0.770128 | 0.045765 | 0.404879 |
| 50 | 0.913915 | 0.010614 | 0.153476 |
| 100 | 0.535670 | 0.130372 | 0.228703 |
| 200 | 0.633842 | 0.098714 | 0.476667 |

**Table S2.** *p* values of comparison between ICI and AAC-treated cell viabilities of cell groups in Figure 2.

| single-ICI vs. | PD-L1 | | | | CTLA-4 | | | |
|---|---|---|---|---|---|---|---|---|
| Concentration (nM) | AAC500 | AAC700 | AAC800 | AAC1000 | AAC500 | AAC700 | AAC800 | AAC1000 |
| 3.125 | 0.649564 | 0.049756 | 0.001788 | 0.003817 | 0.782367 | 0.396465 | 0.010784 | 0.009351 |
| 6.25 | 0.793686 | 0.006296 | 0.000033 | 0.000095 | 0.036455 | 0.001418 | 0.001128 | 0.000194 |
| 12.5 | 0.006083 | 0.000564 | 0.000013 | 0.000013 | 0.003471 | 0.000089 | 0.000043 | 0.000003 |
| 25 | 0.022356 | 0.001168 | <0.000001 | 0.000021 | 0.208612 | 0.003033 | 0.000055 | 0.000107 |
| 50 | 0.011244 | 0.001842 | <0.000001 | 0.000001 | 0.158706 | 0.000159 | 0.000069 | 0.000011 |
| 100 | 0.002686 | 0.000837 | <0.000001 | 0.000002 | 0.006308 | 0.000025 | 0.000001 | <0.000001 |
| 200 | 0.000796 | 0.000166 | 0.000002 | 0.000004 | 0.002981 | 0.000023 | <0.000001 | <0.000001 |

| combo-ICIs vs. | PD-L1:CTLA-4 | | | |
|---|---|---|---|---|
| Concentration (nM) | AAC500 | AAC700 | AAC800 | AAC1000 |
| 3.125 | 0.487375 | 0.001129 | 0.000541 | 0.000001 |
| 6.25 | 0.301864 | 0.003203 | 0.000472 | 0.000005 |
| 12.5 | 0.138396 | 0.000006 | 0.000047 | 0.000002 |
| 25 | 0.218936 | 0.000007 | <0.000001 | <0.000001 |
| 50 | 0.262379 | 0.000009 | <0.000001 | <0.000001 |
| 100 | 0.017953 | 0.000054 | 0.000006 | 0.000010 |
| 200 | 0.007163 | 0.000052 | 0.000006 | 0.000008 |

| PS800 vs. | PD-L1 | | CTLA-4 | | PD-L1:CTLA-4 | |
|---|---|---|---|---|---|---|
| Concentration (nM) | single-ICI | mono-AAC800 | single-ICI | mono-AAC800 | combo-ICI | bi-AAC800 |
| 3.125 | 0.253936 | 0.001532 | 0.967623 | 0.075409 | 0.000735 | 0.001222 |
| 6.25 | 0.061533 | 0.000037 | 0.562057 | 0.011179 | 0.001940 | 0.000789 |
| 12.5 | 0.007783 | 0.000015 | 0.210662 | 0.000029 | 0.002459 | 0.000077 |
| 25 | 0.082146 | <0.000001 | 0.848384 | 0.000059 | 0.000006 | <0.000001 |
| 50 | 0.002435 | <0.000001 | 0.826528 | 0.000156 | 0.000045 | <0.000001 |
| 100 | 0.000326 | <0.000001 | 0.004371 | 0.000002 | 0.000989 | 0.000002 |
| 200 | 0.000639 | <0.000001 | 0.111825 | 0.000215 | 0.000834 | <0.000001 |

**Table S3.** *p* values of comparison between the cell viabilities of two differently treated cell groups in Figure 3.

| Treatment | Day 0 | Day 4 | Day 8 | Day 12 | Day 16 | Day 20 |
|---|---|---|---|---|---|---|
| saline vs. nAAC | 0.496155 | 0.4514073 | 0.009644 | 0.246835 | 0.456002 | 0.764895 |
| nAAC vs. single-ICI | 0.054561 | <0.000001 | 0.181792 | 0.504864 | 0.003432 | 0.000263 |
| nAAC vs. mono-AAC | 0.233337 | <0.000001 | 0.583466 | 0.000003 | 0.000012 | 0.000141 |
| mono-AAC vs. bi-AAC | 0.694958 | 0.000055 | 0.002848 | 0.544421 | 0.031382 | 0.040706 |
| combo-ICIs vs. bi-AAC | 0.881679 | 0.000008 | 0.002122 | 0.780589 | 0.173412 | 0.066989 |

**Table S4.** *p* values of comparison between the bioluminescence intensities of two differently treated mouse groups on each day in Figure 4c.
